# A Convolutional Deep Learning Approach to identify DNA Sequences for Gene Prediction

**DOI:** 10.1101/2025.10.03.680292

**Authors:** Jesus Antonio Motta, Pedro David Gomez

## Abstract

In this work, we present a highly efficient machine learning method for identifying DNA sequences that code for genes. The learning process is based on Human Genome Build 38 (GRCh38) sequences extracted from various specialized databases. The DNA sequences were then translated into amino acid sequences and used to build matrices that facilitate the extraction of features with the TF×IDF vectorization for the creation of the training space. The prediction functions are learned using a convolutional neural network (CNN) deep learning model. The training spaces were created using the 24 chromosomes of the human genome and approximately 36,000 genes and pseudogenes whose names were fetched from the HUGO Gene Nomenclature Committee (HGNC). Performance analysis was performed on 24 genes associated with genetic disorders, as well as the surrounding DNA regions. The metrics used were precision, recall, F_score measure, accuracy and ROC curves for the genes of interest. The results achieved exceed all our expectations and place the work at the level of the state of the art for gene prediction.

## 1. INTRODUCTION

THE development of efficient models for gene prediction is a task that presents numerous challenges as well as the development of complex processes that generally consume large resources. It is therefore necessary to undertake certain actions to address problems such as the analysis of very large genomic sequences to identify functional elements such as protein-coding regions and their regulatory regions. For example, in a eukaryotic organism, different situations arise: numerous non-coding regions (introns), a single gene can produce multiple proteins (alternative splicing), several repeated sequences, and the fact that genes evolve differently depending on the species or human groups (allelic frequencies) among others. Let’s think for example about some species that live in different habitats or other organisms, they are subject to selective processes that necessarily determine the evolution of a gene (evolutionary genomics) over time [1], [2], [3].

After the complete sequencing of the human genome was completed in 2003 [4], gene prediction was significantly enhanced, primarily by having a complete genome annotation and the introduction of new techniques based on machine learning. More recently, the use of deep learning-based algorithms running on GPUs has been a key factor in the development of gene prediction.

In this work, we present an efficient method to predict genes from a DNA sequence using the Convolutional Neural Networks learning method and matrices created with the TF**×**IDF metric from Amino-Acids sequences.

## 2. SOME METHODS AND TECHNIQUES FOR GENE PREDICTION

In this part of our work, some developments (in the area of gene prediction) are presented, according to a classification that is related to the way in which DNA sequences are approached and treated for processing.

### 2.1. AB INITIO Methods

The methods based on this approach are characterized by the fact that their analysis focuses solely on the DNA sequence itself. Below we mention some important tools based in this approach that are currently used:

#### 2.1.1 GENSCAN

Is a tool developed at Stanford University [5], [6], that uses Generalized Hidden Markov Model (GHMM) to identify genes. Other important works that describe and demonstrate the use of HMM for gene prediction can be found in [7], [8].

#### 2.1.2 AUGUSTUS

Is used to identify genes in **eukaryotic genomic sequences**. It is trained on over 100 species [9], [10]. It is possible to improve predictions by adding RNA-Seq data and protein alignments.

#### 2.1.3 GeneMark

Used to identify protein-coding regions in DNA sequences (genes or gene parts). To analyze the sequences, it uses statistical methods such as Inhomogeneous Three-Periodic Markov Models [11], [12]. This method has been widely used in both prokaryotic genomes (bacteria, archaea) and eukaryotic genomes (plants, animals, fungi), as well as in metagenomes (mixed microbial communities) and transcriptomes (RNA sequences).

### 2.2. Similarity Based

This approach works by aligning an unknown DNA sequence with other known gene or protein sequences that are retrieved from large databases. If there is a good alignment (similarity), the gene or protein is marked as a candidate gene.

Some methods based on this approach are:

#### 2.2.1 BLAST (Basic Local Alignment Search Tool)

Uses heuristic algorithms to find similarities between an input sequence (DNA, RNA, or proteins) and other sequences in a database. It is one of the best-known tools of this genre [13], [14]

#### 2.2.2 Exonerate

This method has two ways to traverse the state space: heuristic and exhaustive. It is a good tool to identify genes and its locations by aligning **cDNA or mRNA sequences**, Protein sequences and **DNA to DNA sequences** [15]

#### 2.2.3 PhyloCSF (Phylogenetic Codon Substitution Frequencies)

This method identifies coded and non-coded regions in a DNA sequence, comparing multiple sequences from genomes of different species and analyzing the probability that a sequence has evolved toward a coded or non-coded region [16]. Although this method is not exactly a tool for predicting a gene from a biological sequence (DNA, RNA, etc.), it is a very effective aid in the task of annotating non-coding functional elements. Thus, it could be of great help in the investigation of key regulatory elements of the genome.

### 2.3. Unsupervised Machine Learning

In this category we will only mention clustering-based algorithms, taking into account that almost all of the algorithms mentioned in the previous categories use unsupervised methods for their prediction kernels. For example, GenMark uses Markov models, GenScan uses Hidden Markov Models (HMM), Augustus is based on GHMM (Generalized Hidden Markov Models), and others.

#### 2.3.1 Clustering

Clustering groups similar DNA or RNA sequences together based on shared patterns. Some of the main types of clustering used are: Hierarchical Clustering [17], which builds a tree of clusters; K-means [18], which partitions data into k similar clusters; and Alignment-Free Method, which creates clusters from k-mer frequencies [19].

### 2.4. Supervised Machine Learning

In this section of the study, we present the most recent works based on supervised machine learning to predict genes:

In [20], different methods are compared: *regularized regression, instance-based, ensemble* and *deep learning* methods. It concludes that their performance is very similar for the particular data set used. In the work presented in [21], a model for the prediction of gene expression is built, using methylation data. It utilizes **adaptive convolutional layers** [22] and **residual blocks** [23] to handle high-dimensional methylation. In [24], a review of supervised and unsupervised learning methods and deep learning is presented to infer gene networks from genes, proteins, or metabolites (molecules that are produced or used during metabolism). Spectral clustering data (eigenvalues and eigenvectors are extracted from data) is used in [25] for feature extraction and to build a **learning model** to predict gene-function associations. in [26], a method (CNN-MGP) is developed for classify DNA fragments as coding vs. non-coding in metagenomic data, using CNN to automatically learn sequence patterns that distinguish genes from background DNA, avoiding handcrafted features. It achieves a accuracy of 0.91. Prediction whether a specific enhancer interacts with a promoter method (DeepEPI) is presented in [27] combining CNN’s and sequence encodings (OneHot, DNA2Vec) to learn regulatory sequence patterns that drive long-range gene regulation. Its precision is around 0.90. Another enhancer promoter (EPI-DynFusion) is presented in [28] that integrates CNN’s, Transformers, and Bi-GRU’s with an attention-based fusion mechanism to capture both local motifs and long-range dependencies, it achieves an accuracy of 0.83 for AUPR and 0.96 for AUROC. CNNs is used in [29] to learn motif-like patterns directly from sequence, replacing traditional PWM-based motif discovery achieving a performance around 0.90 (DeepBind). An accuracy between 0.89 and 0.95 is presented in [30] with a CNN trained on massive ENCODE datasets to learn regulatory grammar directly from a sequence (DeepSEA). In [31] a method (Basset) is developed using Multi-task CNN that learns shared regulatory features across many cell types simultaneously achieving a precision between 0.90 and 0.95. A method (DanQ) that combines CNNs (motif detection) with bidirectional LSTMs (long-range dependencies), capturing both local and global sequence structure is presented in [32]. This method achieves an accuracy around 0.96.

More recent models have pushed beyond local motif detection toward capturing long-range genomic interactions. **Enformer** [33] represents a major leap in this direction, using transformer-based attention mechanisms to model up to 200 kilobases of sequence context. This enables accurate prediction of gene expression and chromatin features, with correlations often in the 0.60–0.80 range. Similarly, **Basenji2** [34], **Sei** [35], and **ChromBPNet** [36]refine multi-task regulatory prediction with improved architectures, bias correction, and interpretable motif discovery. These models consistently achieve high AUROC and AUPR values, often exceeding 0.90 across diverse regulatory tasks.

In parallel, transformer-based sequence models such as **DNABERT-2** [37] have emerged, applying large-scale pretraining on genomic k-mers to achieve strong performance on enhancer, promoter, and TF-binding classification tasks. DNABERT-2 typically reports accuracy values above 90% and competitive AUROC scores, demonstrating the power of language-modeling approaches when applied to DNA.

## 3. DESIGN AND CONSTRUCTION OF THE MODEL

For the design, implementation, and construction of the prediction functions, all 24 chromosomes of the human genome and a total of approximately 36,000 genes and pseudogenes were considered. The genomic sequences corresponding to these genes and pseudogenes were fetched from the hg38 (GRCh38) builds [38], which are resident in the databases maintained by NCBI [39], Ensembl [40], UCSC [41] and Uniprot [42]. In our study, we were particularly interested in 24 genes (from hundreds of cases) linked to certain diseases due to mutations in the DNA sequence (single gene mutation). In Tables 1, 2 and 3, we present the list of these genes, which includes a brief description of their function, the effect of the mutation and the chromosome to which they belong. As we have said, the names of the genes (approved symbols) are fetched from HGNC (HUGO Gene Nomenclature Committee) [4],[43], which is the entity responsible for assigning symbols and names to human genes. In Table 5, we show the performance of predicting genes from chromosomes and segments (partitions) that contain these genes.

**Table 1.** Interest genes (Genetic disorders)

| Gene | Chromosome | Gral. Function | Mutation effect |
| --- | --- | --- | --- |
| 1. HTT[39], [52] | 4 | Encodes the **huntingtin muscle protein**, which plays a crucial role in nerve cells, as cell **signaling, protein interactions, and DNA repair** | Huntington's disease, a neurodegenerative disorder |
| 2. MYBPC3[53], [54] | 4 | Encodes **cardiac myosin-binding protein C (cMyBP-C)**, a **structural and regulatory protein in the heart's contractile function** | Left ventricular hypertrophy. Arrhythmias, sudden cardiac death. Ventricular dilation, systolic disfunction, stiff ventricles |
| 3. HLA-DRB1 [52], [53] | 6 | Plays a role in **autoimmune diseases**, like **rheumatoid arthritis, multiple sclerosis, and autoimmune Addison disease** | Affects immune responses differently. Par example, specific variants are associated with **autoimmune Addison disease**, where the immune system mistakenly attacks the adrenal glands |
| 4. COL2A1 [39], [52], [53] | 12 | A very important protein for the formation of **cartilage, joints, eyes, and the inner ea** | **Stickler syndrome (affecting vision, hearing).** **Spondyloepiphyseal dysplasia congenita (SEDC) (a skeletal disorder).** **Kniest dysplasia (leading to short stature and joint problems).** **Achondrogenesis type 2 (a severe skeletal disorder)** |
| 5. BRCA2 [39], [52] | 13 | Is a **tumor suppressor**. Plays a crucial role in **DNA repair**. | Increase the risk of **several cancers**, most notably **breast and ovarian cancer**, but also **prostate and pancreatic cancer** |
| 6. APOE [39], [52], [53] | 19 | A protein that helps transport cholesterol and other fats through the bloodstream | **The variant APOE ε4** increases the risk of developing **Alzheimer's disease** and is linked to more severe forms of the condition |
| 7. PTPN2 [52], [53] | 1 | Encodes **protein tyrosine phosphatase non-receptor type 22**, which plays a essential role in **immune system regulation** | Have been linked to an increased risk of **autoimmune diseases**, including:<br>**Type 1 diabetes, Rheumatoid arthritis, Systemic lupus erythematosus, Graves' disease, Hashimoto's thyroiditis** |
| 8. FTO [39], [52], [53] | 16 | Is linked to **body weight regulation and metabolism** | Variants of the **FTO gene** have been associated mainly with **higher body mass index (BMI) and obesity risk** |
| 9. LEPR [39], [52], [53] | 1 | Encodes the **leptin receptor**, a protein that plays a crucial role in **appetite regulation, metabolism, and energy balance** | **Leptin receptor deficiency**, which is associated with **severe obesity, excessive hunger, and hormonal imbalances** |
| 10. HFE[39], [52], [53] | 6 | Plays a definitive role in **iron metabolism** by regulating how the body absorbs iron from food | **Hereditary hemochromatosis**, where excess iron builds up in organs like the **liver, heart, and pancreas**, potentially leading to serious health issues such as **liver disease, diabetes, and heart problems** |
| 11. LDR [52], [53] | 19 | Produces a lipoprotein receptor (LDLR) that helps remove **LDL (bad cholesterol)** from the bloodstream by binding to LDL particles and transporting them into cells for processing | **Familial hypercholesterolemia (FH)**: leads to **high cholesterol levels**, increasing the risk of **heart disease and early-onset heart attacks** |

**Table 2.** Interest genes (Genetic disorders)

| Gene | Chromosome | Gral. Function | Mutation effect |
| --- | --- | --- | --- |
| 12. CFTR [52], [53] | 7 | Encodes a **chloride ion channel** that plays a crucial role in maintaining the balance of salt and water on cell surfaces, particularly in the **lungs, pancreas, and digestive system** | **Cystic fibrosis (CF)**, that causes thick, sticky mucus to build up in the lungs and other organs |
| 13. FBN1[52], [53] | 15 | Encodes **fibrillin-1**, a very important protein in the **extracellular matrix** that helps form **microfibrils**, which provide structural support to tissues like **skin, blood vessels, and bones** | **Marfan syndrome**, affecting the **heart, eyes, and skeleton**.<br>**Weill-Marchesani syndrome**, which leads to **short stature and joint stiffness**.<br>**Ectopia lentis**, where the **lens of the eye becomes displaced**, affecting vision |
| 14. HBB [52], [53] | 11 | Encodes **hemoglobin subunit beta**, a essential component of hemoglobin—the protein responsible for carrying oxygen in red blood cells | **Sickle Cell Disease**: leads to abnormal hemoglobin that distorts red blood cells into a sickle shape, causing **pain, anemia, and organ damage** |
| 15. FMR1 [52], [53] | X | Plays a fundamental role in **brain development and cognitive function**. It encodes the **FMRP protein**, which helps regulate **synaptic plasticity**, influencing learning and memory | **Fragile X Syndrome (FXS)**: Caused by an expansion of the **CGG nucleotide repeats**, leading to intellectual disability and developmental delays<br>**Fragile X-associated primary ovarian insufficiency (FXPOI)**: Affects ovarian function and fertility<br>**Fragile X-associated tremor/ataxia syndrome (FXTAS)**: A neurodegenerative disorder |
| 16. HBA1 [53], [54] | 16 | **HBA1** encodes **hemoglobin subunit alpha 1**, a key component of hemoglobin—the protein responsible for carrying oxygen in red blood cells<br>Works with **HBA2**, gene that produces alpha-globin | **Alpha Thalassemia**: Caused by deletions or mutations in **HBA1 and HBA2**, leading to reduced hemoglobin production<br>**Hemoglobin H Disease**: Results from the loss of three alpha-globin alleles, causing **anemia, jaundice, and enlarged spleen**<br>**Hb Bart Syndrome**: The most severe form, leads to **hydrops fetalis** (severe fluid buildup before birth) |
| 17. HBA2 [53], [54] | 16 | Encodes **hemoglobin subunit alpha 2**, which is a key component of hemoglobin—the protein responsible for carrying oxygen in red blood cells | **Alpha Thalassemia**<br>**Hemoglobin H Disease**<br>**Hb Bart Syndrome** |
| 18. BRCA1 [39], [52] | 17 | Is a **tumor suppressor gene** that plays a vital role in **DNA repair** and maintaining genomic stability | Mutations in **BRCA1** significantly increase the risk of **breast and ovarian cancer**, as well as other cancers like **pancreatic and prostate cancer** |
| 19. ACE[52], [54] | 17 | Encodes the angiotensin-converting enzyme, which is a very important regulator of blood pressure, fluid balance, and electrolyte homeostasis | Impaired kidney development, low blood pressure, anuria, hypertension, stroke, diabetic nephropathy, late-onset Alzheimer's disease |

**Table 3.** Interest genes (Genetic disorders)

| Gene | Chromosome | Gral. Function | Mutation effect |
| --- | --- | --- | --- |
| 20. COMT | 22 | Enzyme that breaks down catecholamine neurotransmitters (dopamine, norepinephrine, epinephrine). Important for cognition and stress response. | Alters dopamine metabolism; affects cognition, pain sensitivity, and risk modulation for psychiatric traits. |
| 21. FOXP2 | 7 | Transcription factor essential for neural circuits involved in speech and language. | Causes developmental verbal dyspraxia; impaired speech motor control and language processing. |
| 22. DMD | X | Produces dystrophin, a structural protein that stabilizes muscle cell membranes. | Duchenne or Becker muscular dystrophy; progressive muscle weakness and cardiomyopathy |
| 23. SRY | Y | Master regulator of male sex determination; initiates testis development. | Disorders of sex development: XY gonadal dysgenesis or XX testicular development depending on mutation/translocation. |
| 24. HNF1B | 17 | Transcription factor for kidney, pancreas, liver, and urogenital development. | MODY5 diabetes, kidney cysts/malformations, electrolyte abnormalities, pancreatic hypoplasia |

We have divided the design and construction process of our model into the following 7 stages: *DNA sequence fetching, cleaning, partitioning, feature engineering (rebuilding features), training and test set selection, function induction and performance measurement*.

### 3.1 Sequence Fetching

The DNA sequences that form the training space for the learning functions are fetched from reputable specialized databases. We have taken approximately 36,000 gene-coding sequences from each of the 24 chromosomes of the human genome from a list provided by HGNC [43]. The accessed databases were NCBI [38], Ensembl [40], and UCSC [41]

### 3.2 Cleaning

Sequences fetched from databases undergo a cleansing process. It begins by verifying their existence and format, as well as the standardization of upper or lower case. Then, blanks, special characters, numeric characters, and ambiguous bases are removed (only A, T, G, and C are considered).

### 3.3 Partitioning

Considering the size of the training space (DNA sequences of 24 chromosomes containing code for approximately 36,000 genes and pseudogenes) and to facilitate efficiency in the different tasks involved, we have decided to use a “divide and conquer” approach: each chromosome *i* is divided into *j* partitions containing its corresponding genes g_i,j._ Thus, for example, chromosome 1 (which is the largest human chromosome), containing approximately 3500 genes and pseudogenes, has been divided into 12 partitions of approximately 292 genes each. Thus, if **nc** = number of chromosomes and **np** = number of partitions, then the total number of genes in the training space is:

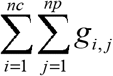

### 3.4 Feature Engineering

The feature engineering stage consists of the following sub-stages:

#### 3.4.1 ORF Identification

An ORF (*Open Reading Frame*) is a stretch of DNA that has the potential to create a protein. This sequence always begins with the ATG (methionine) codon and ends with one of these three codons: TAG, TGA, TAA. The National Human Genome Research Institute (NHGRI) defines an ORF as a stretch of DNA that when transcribed into RNA, has no stop codon [44]

Considering that the presence of ORFs in a DNA sequence could indicate the existence of a gene (because it contains coding for amino acids), in our research, we will use ORFs in a different approach. It is not strictly a matter of identifying the presence of a gene in the DNA sequence. We have fetched the genes of each chromosome based on the exact chromosomal location, a single accession ID, or by indicating the range of its location in the DNA sequence that contains it, ensuring that the direction is 5’ to 3’.

To identify the ORFs, the following steps are performed:

##### a) Identification/Extraction of Codons (triplets of nucleotides)

All DNA-based genomes contain 64 types of triplets (codons), created from 4 nucleotides A, T, G, C (4^3^ = 64). This would result in 61 codons encoding 20 amino acids and 3 codons encoding stop signals (TAG, TGA, TAA). Each gene sequence is examined, and its codons are identified.

##### b) Building Reading Frames

The codon sequence is converted into 6 different reading frames (rf): 3 on the forward strand (+1, +2, +3), and 3 on the reverse complement (-1, -2, -3) as shown below: Starting for example with the sequence: ‘ATGCGTACGTAGCTAGCTAAATGCGTGAATGCGTACGTAGCTAGCTAAATGCGTGA’, then: **Frame 1**: is the starting sequence with its identified codons: [‘ATG’, ‘CGT’, ‘ACG’, ‘TAG’, ‘CTA’, ‘GCT’, ‘AAA’, ‘TGC’, ‘GTG’, ‘AAT’, ‘GCG’, ‘TAC’, ‘GTA’, ‘GCT’, ‘AGC’, ‘TAA’, ‘ATG’, ‘CGT’]

**Frame 2**: is formed from the second nucleotide, leaving the first and then grouping again 3 by 3: [‘TGC’, ‘GTA’, ‘CGT’, ‘AGC’, ‘TAG’, ‘CTA’, ‘AAT’, ‘GCG’, ‘TGA’, ‘ATG’, ‘CGT’, ‘ACG’, ‘TAG’, ‘CTA’, ‘GCT’, ‘AAA’, ‘TGC’, ‘GTG’]

**Frame 3**: is formed from the third nucleotide, leaving the first 2 and then grouping again 3 by 3: [‘GCG’, ‘TAC’, ‘GTA’, ‘GCT’, ‘AGC’, ‘TAA’, ‘ATG’, ‘CGT’, ‘GAA’, ‘TGC’, ‘GTA’, ‘CGT’, ‘AGC’, ‘TAG’, ‘CTA’, ‘AAT’, ‘GCG’, ‘TGA’

**Reverse complement:** TCACGCATTTAGCTAGCTACGTACGCATTCACGCATTTAGCTAGCTACGTACGCAT’ Now, the same processes of frames 1, 2 and 3 are applied:

**Frame -1:** [‘TCA’, ‘CGC’, ‘ATT’, ‘TAG’, ‘CTA’, ‘GCT’, ‘ACG’, ‘TAC’, GCA’, ‘TTC’, ‘ACG’, ‘CAT’, ‘TTA’, ‘GCT’, ‘AGC’, ‘TAC’, ‘GTA’, ‘CGC’]

**Frame -2:** [‘CAC’, ‘GCA’, ‘TTT’, ‘AGC’, ‘TAG’, ‘CTA’, ‘CGT’, ‘ACG’, ‘CAT’, ‘TCA’, ‘CGC’, ‘ATT’, ‘TAG’, ‘CTA’, ‘GCT’, ‘ACG’, ‘TAC’, ‘GCA’]

**Frame -3:** [‘ACG’, ‘CAT’, ‘TTA’, ‘GCT’, ‘AGC’, ‘TAC’, ‘GTA’, ‘CGC’, ‘ATT’, ‘CAC’, ‘GCA’, ‘TTT’, ‘AGC’, ‘TAG’, ‘CTA’, ‘CGT’, ‘ACG’, ‘CAT’]

##### c) Identification of the ORF’s

The sequences in the frames are examined to identify those that begin with ATG and end with one of the three termination signals TAG, TGA, TAA. Thus, we find an ORF in Frames 1 and 3(highlighted)

#### 3.4.2 Conversion of ORF stretches to amino acid sequences

The conversion of ORFs to amino acid sequences occurs when the nucleotide T (thymine) is converted into U (uracil) in the physiochemical process of the cell that passes from a DNA sequence to an RNA sequence through the action of the enzyme RNA polymerase.

If we now take the ORF sequences from frames 1 and 3 above, then:

ORF (Frame 1): [‘ATG’, ‘CGT’, ‘ACG’, ‘TAG’ → AUG CGU ACG UAG →*met arg thr*

ORF (Frame 3): [‘ATG’, ‘CGT’, ‘GAA’, ‘TGC’, ‘GTA’, ‘CGT’, ‘AGC’, ‘TAG’] →AUG CGU GAA UGC GUA CGU AGC →*met arg glu cys val arg ser*

The transfer of an initial DNA sequence to another amino acid sequence in the Training Set (TS) construction pathway to obtain the prediction functions, allows us, firstly, to clearly differentiate *exons* from *introns*, thus enhancing the discriminatory power of the coded part of the DNA sequence, that is, the portion that determines its function in protein production. Secondly, redundancy is reduced, taking into account that many codons produce the same amino acid (e.g., CUU and CUA, which code for leucine). This approach allows us to have more expressive sequences with great discriminatory power, a very important requirement for obtaining good prediction functions.

It is important to note that the above processes are performed automatically using powerful functions provided by different libraries. In our case, we used the Scikit-Learn(sklearn) and Bio-Python libraries.

#### 3.4.3 Construction of TF×IDF matrices

**TF×IDF** [45] is a classic method for feature extraction, initially used in text mining and Natural Language Processing (NLP). This weighting scheme identifies words that are frequent in a specific document but not very common in the entire dataset (corpus). This makes it an excellent resource for determining features and representing text in machine learning (ML) for classification, clustering, or information retrieval tasks.

**tf** is calculated as follows:

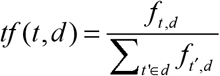

where *f*_*t*,*d*_ is the number of times that term *t occurs* in document *d* and 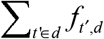 is the total number of terms in the document.

**idf** is calculated with the following formula:

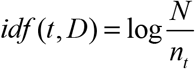

where *D* is the set of all documents, *N = D* is the total of documents and n_t_ is the number of documents where the term *t* occurs.

Then,

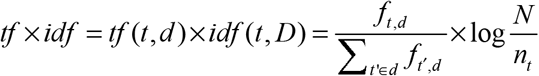

This method has been used to obtain k-mer frequencies (possible DNA or RNA subsequences of length k). An interesting application of this method is presented in the work of [46]. The use of tf×idf-kmer is also found in some other works such as [47], [48] and [49].

In our work, we used the tf×idf method to extract features from amino acid matrices we constructed for each gene on each chromosome. Taking into account that the codon sequences of each gene have already been converted to amino acid sequences, the matrices are constructed as follows: the amino acid strips are taken and 20×20 matrices (corresponding to 20 amino acids that make up proteins) are formed. If any column or row is missing, for example, to construct the final matrix for a given gene, it is completed with the first column or row of the matrix in question. Then, TF-IDF vectors are created by columns of each matrix, thus configuring a TF-IDF matrix.

## 4. TRAINING AND TEST SET SELECTION

The training and test sets are created from the amino acid sequences of each gene on each chromosome partition. The examples are the amino acid chains and their labels are the name of the gene to which the chain corresponds. Each example is a 20×20 TF-IDF matrix. Training sets are created for each P_ij_ partition of each chromosome. The partitions particularly studied contain approximately a total of 23,000 genes and pseudogenes, each corresponding to a “class” within the partition to which it belongs. Table 4 shows the number of examples per chromosome, and Table S2_1 (Supplementary Material #2. Available upon request by contacting the authors) shows the number of examples per chromosome partition. We used an 80/20 approach to split the dataset into training and test sets, as well as a 10% validation set.

**Table 4.**
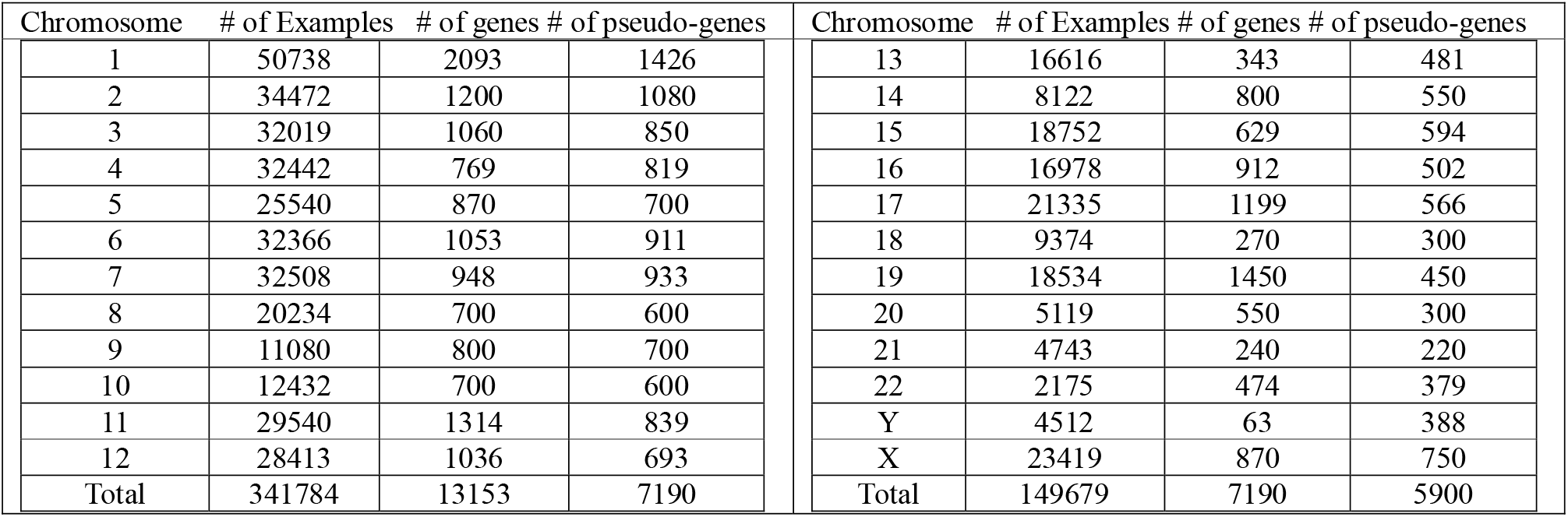
Number of examples/genes/pseudogenes by chromosome.

| Chromosome | # of Examples | # of genes | # of pseudo-genes | Chromosome | # of Examples | # of genes | # of pseudo-genes |
| --- | --- | --- | --- | --- | --- | --- | --- |
| 1 | 50738 | 2093 | 1426 | 13 | 16616 | 343 | 481 |
| 2 | 34472 | 1200 | 1080 | 14 | 8122 | 800 | 550 |
| 3 | 32019 | 1060 | 850 | 15 | 18752 | 629 | 594 |
| 4 | 32442 | 769 | 819 | 16 | 16978 | 912 | 502 |
| 5 | 25540 | 870 | 700 | 17 | 21335 | 1199 | 566 |
| 6 | 32366 | 1053 | 911 | 18 | 9374 | 270 | 300 |
| 7 | 32508 | 948 | 933 | 19 | 18534 | 1450 | 450 |
| 8 | 20234 | 700 | 600 | 20 | 5119 | 550 | 300 |
| 9 | 11080 | 800 | 700 | 21 | 4743 | 240 | 220 |
| 10 | 12432 | 700 | 600 | 22 | 2175 | 474 | 379 |
| 11 | 29540 | 1314 | 839 | Y | 4512 | 63 | 388 |
| 12 | 28413 | 1036 | 693 | X | 23419 | 870 | 750 |
| Total | 341784 | 13153 | 7190 | Total | 149679 | 7190 | 5900 |

### 4.1 Function Induction

As mentioned above, the amino acid sequences of the chromosomes have been divided into partitions containing the gene name (label) and its sequences (examples). Thus, the model will construct prediction functions for each partition of the corresponding chromosome.

Our model’s architecture is based on a CNN (Convolutional Neural Network). This is a type of neural network that uses special layers called convolution layers, due to their similarity to the human brain’s visual cortex (where everything we see is processed), to detect and extract features from input data.

The network architecture is Sequential Conv2D (2-dimensional convolution). The optimization function is **Adam**, and the activation function is **Softmax**. The general parameters used (hyperparameters) were: **Kernels** (filters): 16 (number of patterns to learn), **Kernel size**: 3 (size of the kernel or filter), **Pooling**: Max Pooling (reduction of the input size, but preserving the most relevant information), **Stride:** 1 (step), **Decay rate**: 0.42 (reduction of the learning rate as training progresses) and **Learning rate**: 0.001 (how much the model updates its weights). The layers number was 3.

We used a number of epochs equal to 120, but introduced the parameter **early stopping** equal to 6. Here, we provide a summary of the mathematical foundations of CNN, along with an explanation of how it operates. We will also include a brief application involving the entry [1,1] of the feature map in a tf*idf matrix. We have a tf*idf matrix of size m_1_×m_2_ which will be the tensor T and a kernel K of size n_1_×n_2_. The resulting product will be another array F of dimension (m_1_-n_1_+1) ×(m_2_-n_2_+1).

Then,

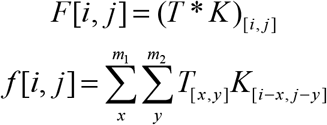

To calculate the [1,1] entry of the future map using the equation provided with an input tf**×**idf matrix of order 20, a kernel of size 3 and a stride of 1, we proceed as follows:

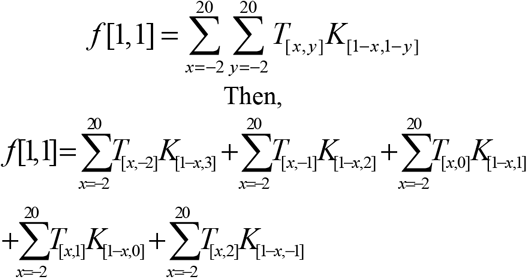

This procedure is repeated for all entries in the matrix.

To address the fact that filters are focused on the center of the matrix, ***padding*** is added by including a column and row of zeros on each side of the tensor.

Then, an ***activation function*** is applied to help the network learn non-linear relationships between matrix features for pattern recognition.

Let *φ*_*a*_ be an activation function and *b* be a bias term, then

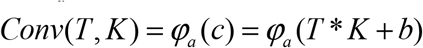

Next, ***pooling P*** is performed. The goal of pooling is to identify the most important features in the convolutional matrix by applying operations that reduce the feature map. We use the Max pooling aggregation function, which selects the maximum value from the feature map.

Let φ _*p*_ be a pooling function and *Conv*(*T, K*) = *C*, then *P* = *φ*_*p*_ (*C*)

After completion the previous steps, the resulting array is flattened into a single vector, which will serve as the input to a fully connected layer for classification using an activation function

Let *g* represents the activation function, *X* be the set of flattened vectors with weights *w*, pooling equal to *P*, and *d* be a bias term, then 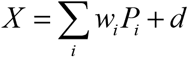 and *z* = *g* (*X*)

Fig. 1 presents a block diagram of the function induction processes, from the input of the examples of a chromosome partition to the output of the predictor functions and their performance values. Fig. 2 presents a more detailed description of the induction process.

**Fig. 1.**
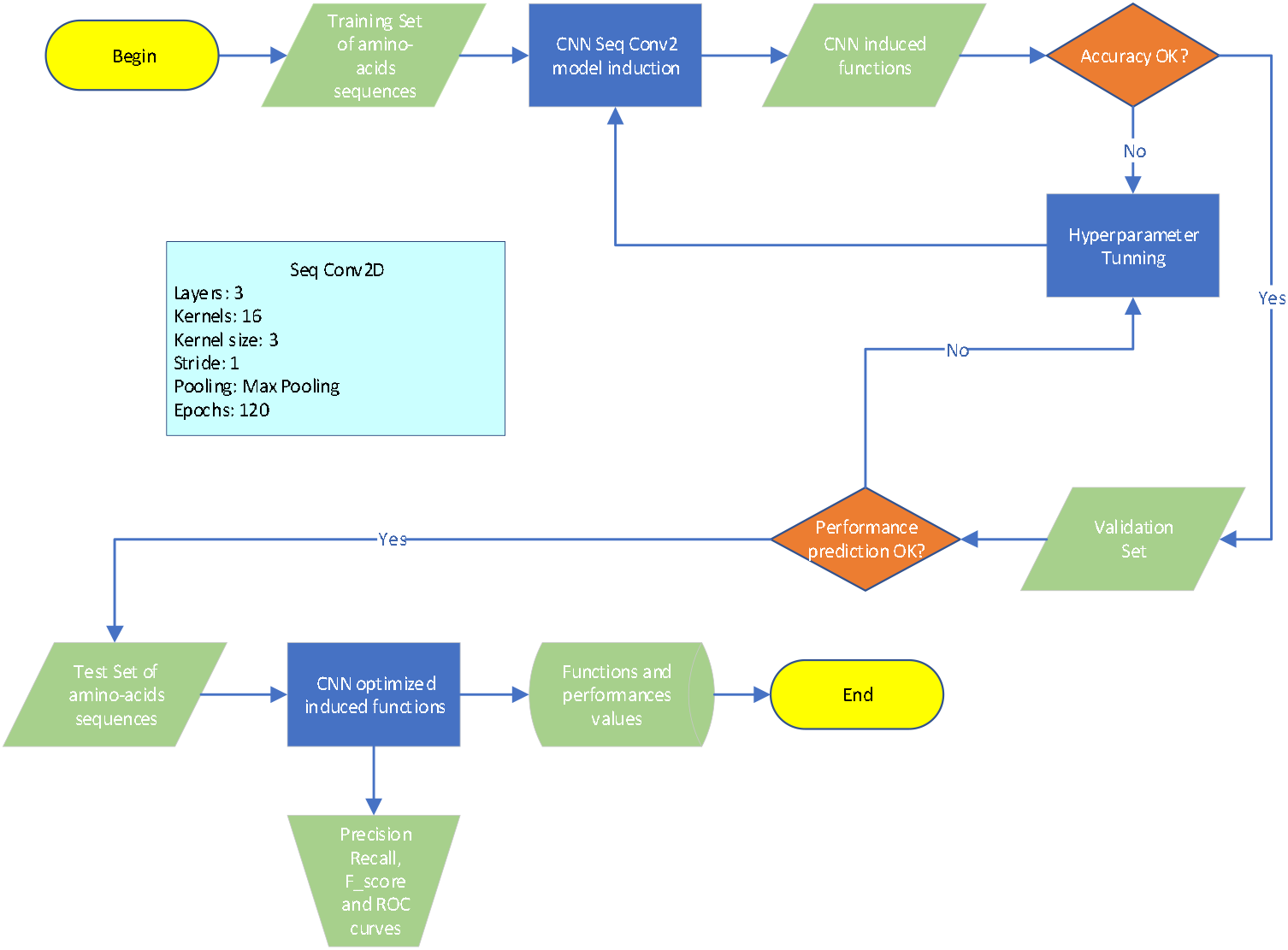
Function induction process

**Figure 2.**
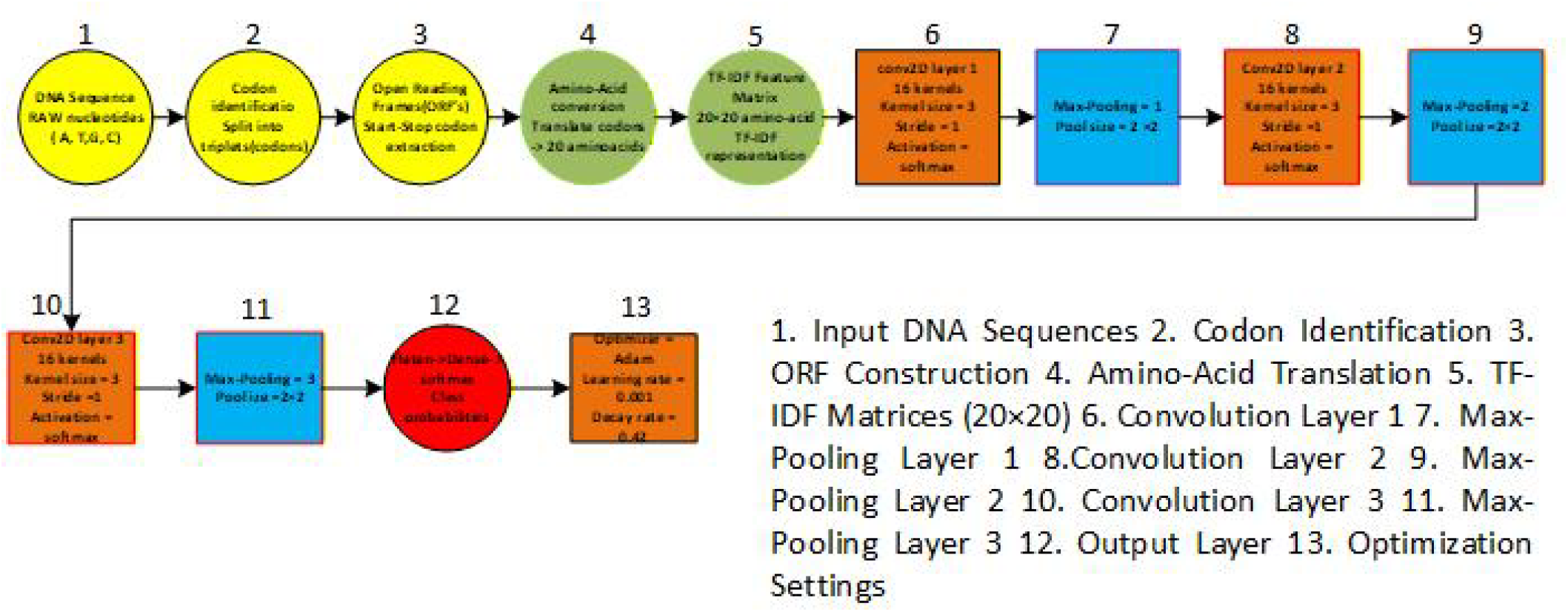
Detailed Induction Processes

Here’s a general description of the hardware and software used:

Hardware: 24 threads Intel CPU, 24 GB GPU card, 64 GB main memory, 6 TB SSD disks Software: Linux Ubuntu 24.04, Python 3.10, Scikit-Learn(sklearn) 1.7.0, TensorFlow 2.19.0, CUDA 11.8, BioPython 1.85

### 4.2 Performance Measurement

To evaluate the performance of our model, we used the Precision, Recall, F__score_ and Accuracy metrics, as well as ROC curves. A description of each is presented below

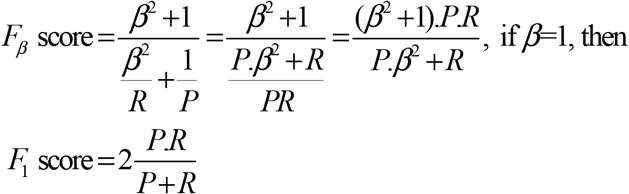

### 4.2.1 Precision(P), Recall(R), F_1_ score and Accuracy (ACC)

Let TP be the number of genes correctly classified by our model, FP the number of genes incorrectly classified, FN are the genes that, despite belonging to the class under study, are classified as not belonging to it and TN and TN which are genes that do not belong to the gene in question. Then, precision (P) recall (R) and accuracy (ACC) are defined as follows:

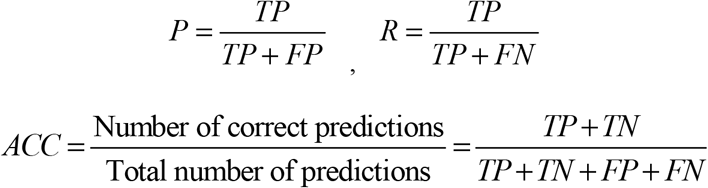

The F_β_-score [50] is the harmonic mean of precision and recall, where β is a constant that represents the importance of recall on precision. It is defined as:

#### 4.2.2 ROC curves[51]

Let N be the number of genes to be classified, TN the number of genes that do not belong to the class of the gene examined, TPR be the rate of genes that belong to the class examined. If TN is the number of genes that do not belong to the class examined, we define the following metrics:

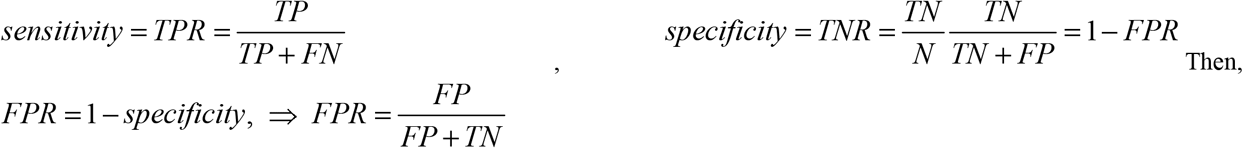

We used **sensitivity** and **specificity** measures to create a **ROC** plot and assess the quality of the prediction by examining the point where the maximum TPR value is obtained and which corresponds to the minimum FPR.

The Area Under the Curve (AUC) of the graph indicates the probability of success of the function to accurately classify a gene vs. the probability of being wrong in this task.

## 5. Discussion

The metrics applied to both, the learning functions to evaluate the model’s ability to fit the data on which it was trained and its ability to learn patterns, as well as the metrics applied to the model’s predictive ability (ability to generalize) on unseen examples from the test set, present excellent results as discussed below.

Table 5 shows the metrics **precision (P), recall (R), F**_**1**_ **score** and **accuracy (ACC)** for each of the partitions containing the genes of interest and on which we focused our attention for the performance evaluation. These partitions correspond to chromosomes 1, 4, 6, 7, 11, 12, 13, 15, 16, 17, 19, 22, X, Y and each one is associated with the predictor function learned from the sequences (examples) they contain.

**Table 5.**
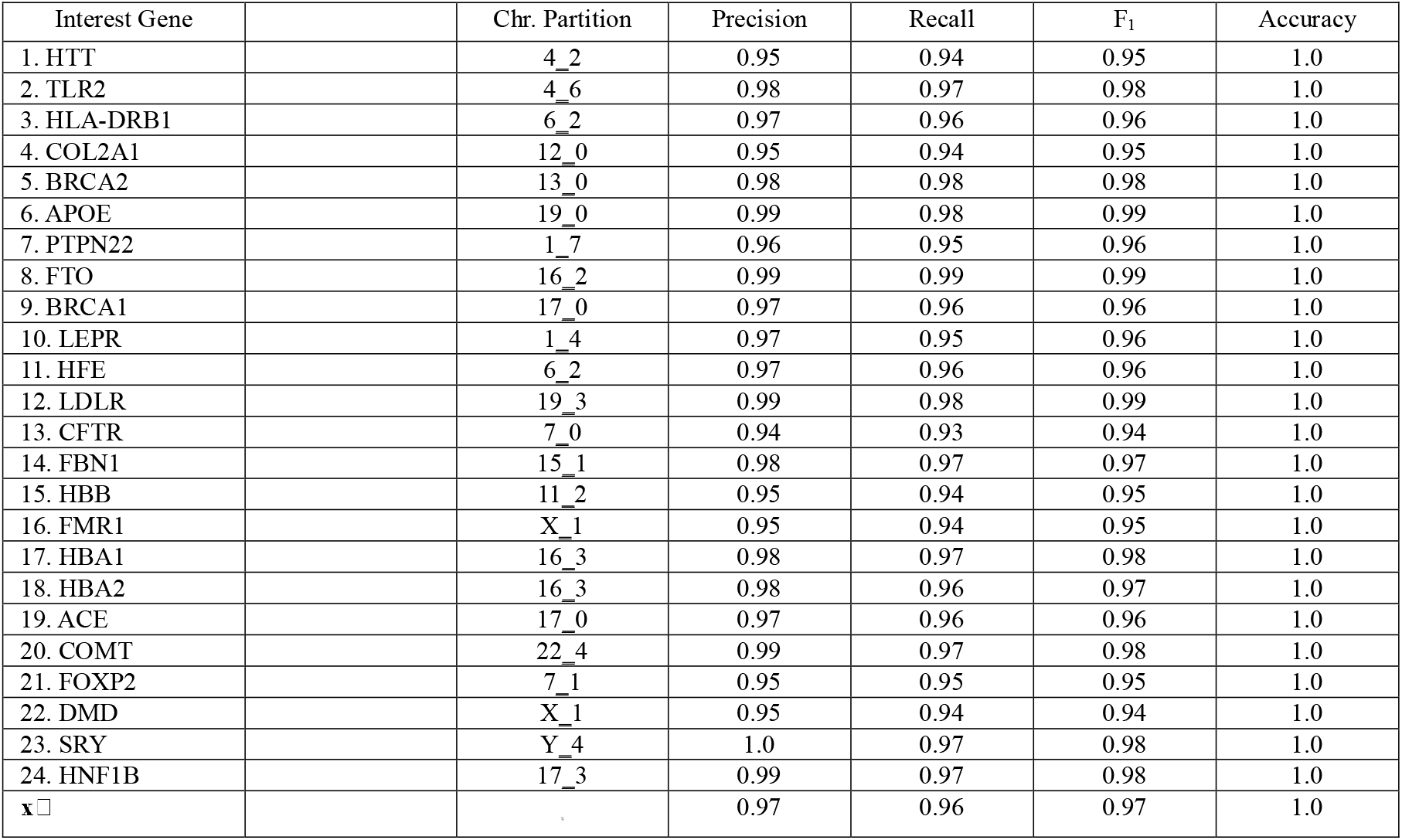
Performance of partitions of genes of interest.

It can be observed that the **Accuracy** for all partitions was equal to 1.0, which indicates the great power of the function to learn the patterns found in DNA (amino acid) sequences and, consequently, their ability to fit these same sequences.

Consistent with these ideas, we found that the **precision (P)** of the gene prediction for chromosome partitions is above 95%, with the sole exception of partition 7_0 (94%), which contains the CFTR gene. We observed that in 67% of cases, this metric was between 97% and 100%. The P value for partitions 16_2, 17_3, 19_3, 22_4, and Y, which contain the HNF1B, SRY, COMT, LDLR, FTO, and APOE genes, was remarkably high, ranging from 99% to 100%. The average across all partitions was 97%. The behavior of **recall (R)** was similar to precision, noting that its average was reduced by one point with respect to this, but at the same time the **F**_**1**_ **score** was significantly high with a value of 97%, which indicates a very good balance between false positives and false negatives of the prediction.

Analyzing the ROC curves presented in figures 3, 4 and figures S1_1 to S1_4(Supplementary Material #1. Available upon request by contacting the authors), we observe that in all of them the maximum value for predictions with “true positive” results corresponds to the minimum value of “false positives”, which indicates the great power of the classifier to truly identify the class (gene) in question. Likewise, we notice that in 71% of cases the AUC value is equal to or greater than 0.95. We also find that in 29% of these values they are between 0.90 and 0.95. Values less than 0.90 were found in only one case, which corresponds to the CFTR gene (0.88). We also observe that the function is extraordinarily efficient in identifying the genes: ACE, APOE, BRCA2, COL2A1, FBN1, HBA1, HBA2, HBB, HFE, HLA-DRB1, HNF1B, LDLR, MYBPC3 and SRY, whose AUC values are equal to 1.0.

**Fig. 3.**
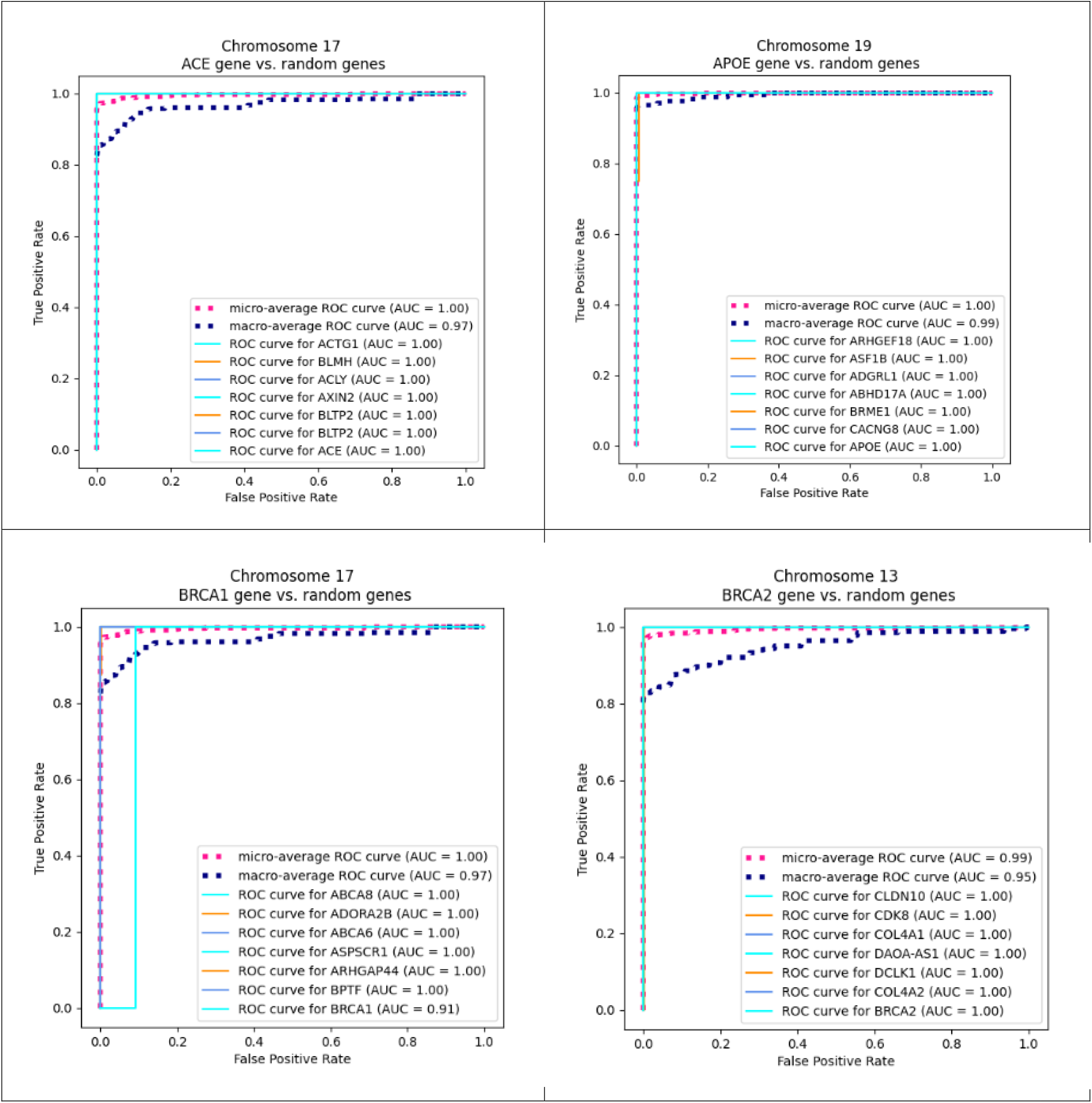
Genes of Interest vs. random genes

**Fig. 4.**
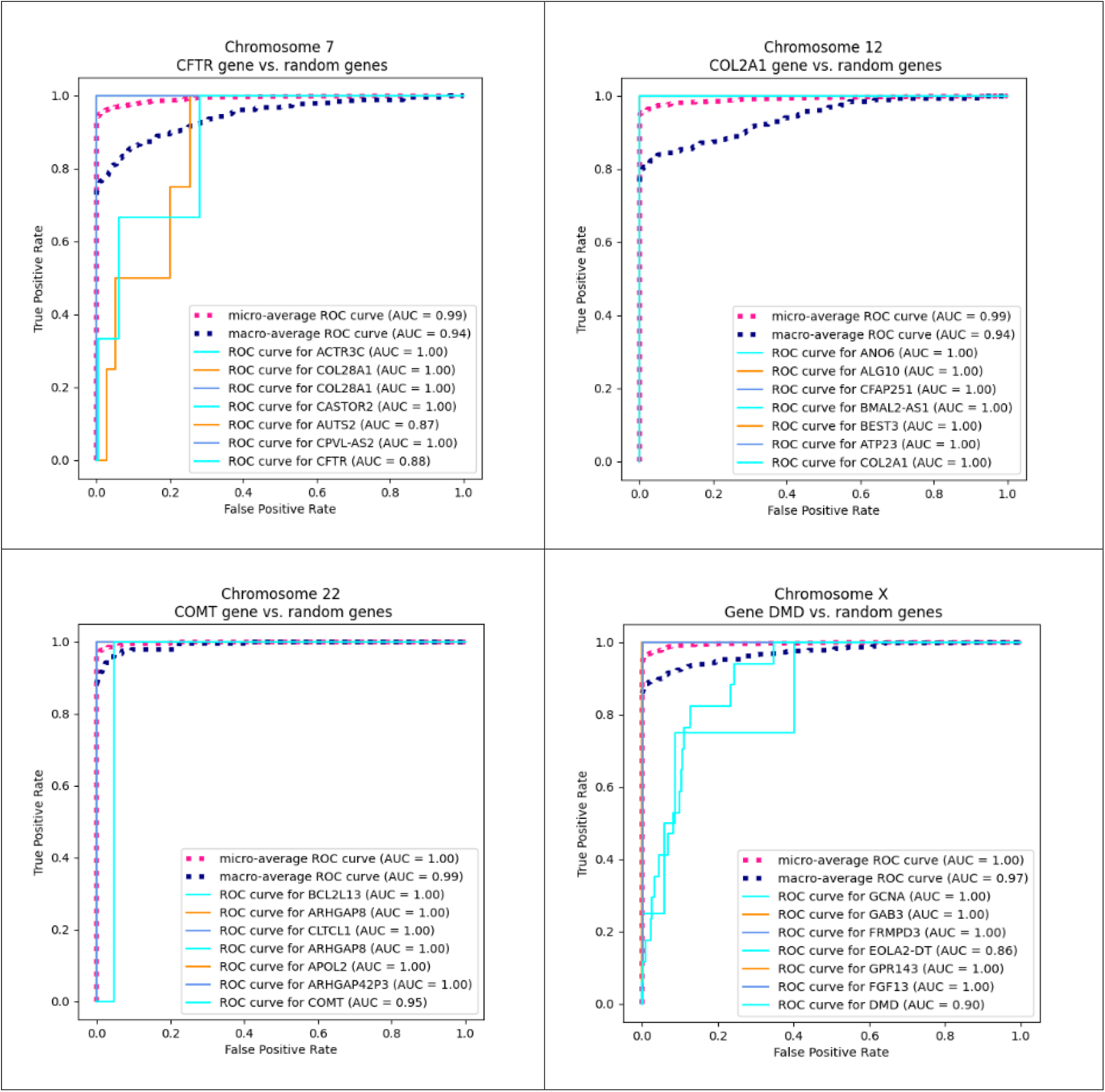
Genes of Interest vs. random genes

Considering that the partitions specially studied contain approximately 23000 genes and pseudogenes, the ROC curves of each of these genes have been constructed by comparing them with 6 other ROC curves of genes randomly extracted from the partition to which the gene belongs, thus also showing the quality of the prediction for additional genes. For example, the ROC curve for the BRCA2 gene on chromosome 13 (see the graph in Fig.3) was compared with the genes CLDN10, CDK8, COL4A1, DAOA-AS1, DCLK1, and COL4A2.

The model achieved perfect classification accuracy (1.0) on the benchmark dataset, with predicted probabilities clustering around 0.98 for coding sequences and below 10□□ for non-coding sequences. This indicates excellent discrimination while maintaining appropriate probabilistic calibration. Let’s keep in mind that accuracy doesn’t care *how confident* the model is. It only cares whether the predicted class matches the true class.

To contextualize the performance of our model, we compared it against AUGUSTUS, a widely used ab initio gene finder based on generalized hidden Markov models. **AUGUSTUS is an ab initio gene prediction system** used to identify protein-coding genes directly from genomic DNA. It is one of the most widely used tools in genome annotation pipelines and is considered the successor to earlier systems like **GENSCAN** and **GeneMark**. We benchmarked our model against this system, which served as a classical HMM-based baseline for gene prediction. It provides an appropriate reference point for evaluating discrimination and robustness

We generated a balanced artificial dataset derived from the 24 protein-coding genes used in training. For each gene, we created four perturbed variants: (i) random point mutations affecting 3% of bases, (ii) codon-preserving shuffles that maintain amino-acid composition while disrupting higher-order structure, (iii) deletion of three consecutive bases, and (iv) insertion of three random bases. The model assigned high probabilities to original coding sequences (median ≈ 0.98) and sharply reduced its scores for all perturbed variants, indicating strong sensitivity to disruptions in coding structure. In contrast, AUGUSTUS, which produces only binary predictions, showed limited sensitivity to several perturbation types, particularly small indels and codon shuffles. These results demonstrate that the model captures fine-grained coding signals and provides calibrated probabilistic outputs that classical HMM-based gene finders cannot offer. The following are some results from the application of baseline comparison metrics:

### Brier score

The model achieved an exceptionally low Brier score (0.0002), indicating near-perfect probabilistic calibration. In contrast, AUGUSTUS obtained a Brier score of 0.7167, reflecting both misclassifications and the inherent limitations of HMM-based gene finders, which output only binary predictions and therefore cannot be calibrated.

### Precision–Recall Performance (Fig. 7)

The model demonstrated excellent discrimination between coding and non-coding sequences, as reflected by a near-perfect precision–recall (PR) curve. True coding sequences consistently ranked above all perturbed and artificial negatives, maintaining high precision even at high recall levels. This indicates that the model not only separates classes effectively but also avoids false positives — a critical property in gene prediction, where mis-classifying non-coding regions as exons can propagate errors through downstream annotation pipelines. In contrast, AUGUSTUS produces a single operating point on the PR plane due to its binary outputs, preventing any meaningful assessment of its ranking ability or threshold-dependent behavior.

### Calibration of Predicted Probabilities (Fig. 6)

Calibration analysis revealed that the model’s predicted probabilities closely matched empirical correctness frequencies. The calibration curve aligned almost perfectly with the diagonal, and the Brier score was extremely low (0.0002), indicating that the model’s confidence estimates are both accurate and reliable. This means that a predicted probability of 0.98 corresponds to an actual ∼98% likelihood of the sequence being coding — a property that is essential for downstream decision-making, threshold tuning, and uncertainty-aware annotation. AUGUSTUS, by design, cannot be calibrated: its predictions are restricted to 0 or 1, resulting in a high Brier score (0.7167) and no interpretable probability structure.

### Divergence between the model and AUGUSTUS (Fig. 5)

**Figure 5.**
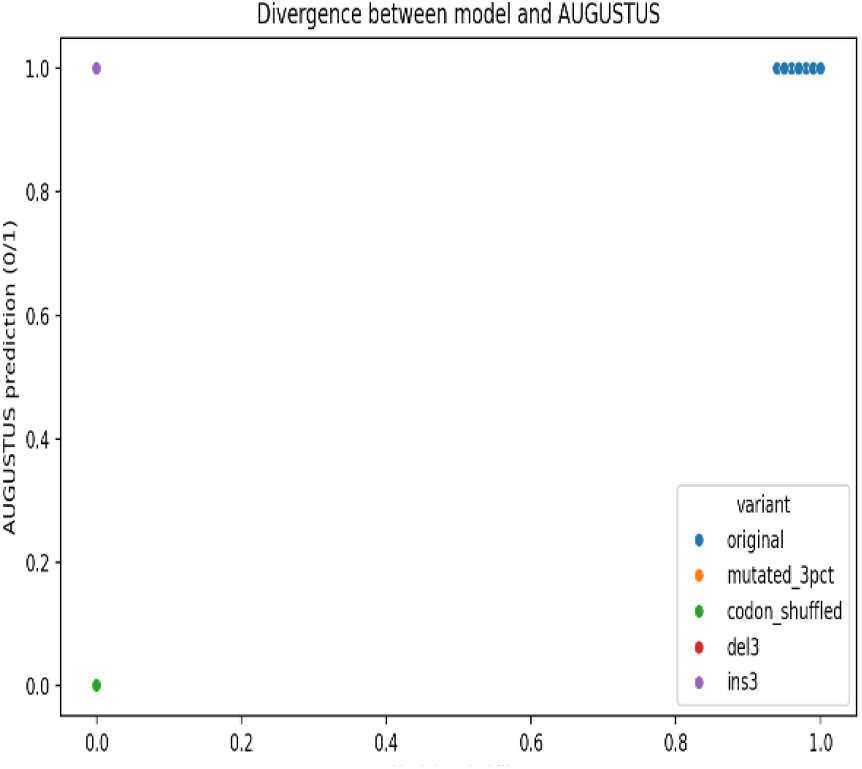
Model’s probab. and AUGUSTUS predictions

Scatterplot comparing the model’s predicted probabilities (x-axis) with AUGUSTUS’s binary coding/non-coding predictions (y-axis) across all perturbation types. Each point represents a sequence variant, colored by perturbation class. AUGUSTUS outputs only discrete labels (0 or 1), forming two horizontal bands, while the model produces continuous probabilities that reflect graded coding potential. Divergence arises where the model assigns low probability to perturbed or disrupted sequences that AUGUSTUS still classifies as coding (upper-left region), and where the model confidently identifies coding sequences that AUGUSTUS fails to detect (lower-right region). These patterns highlight systematic differences between probabilistic sequence modeling and HMM-based gene prediction.

### ROC curve (Fig. 8)

**Figure 6.**
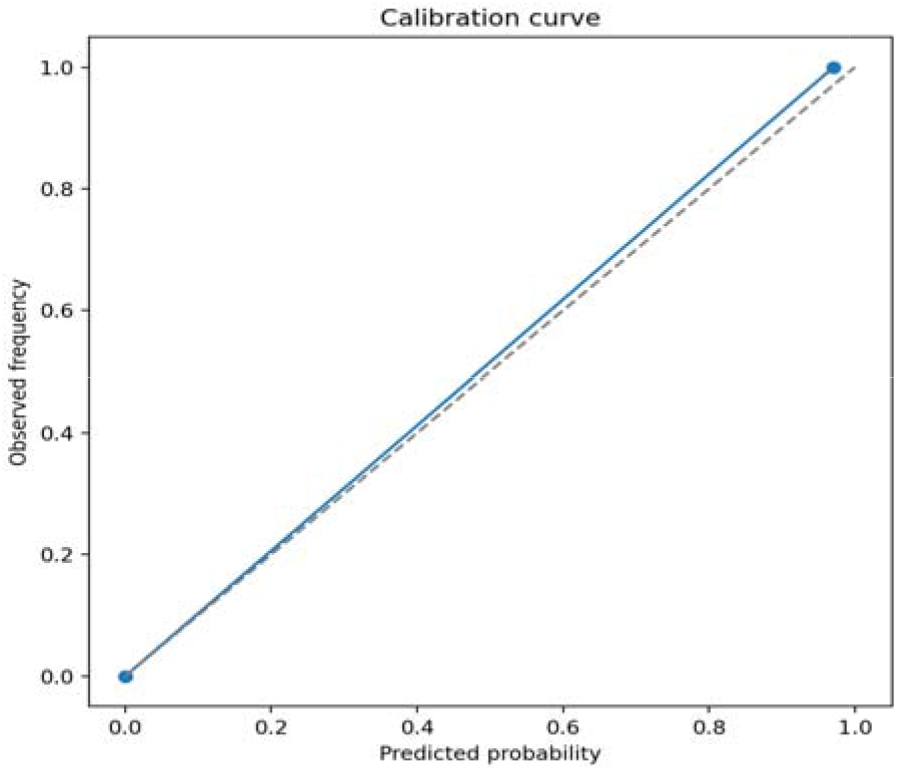
Calibration of prediction probabilities

**Figure 7.**
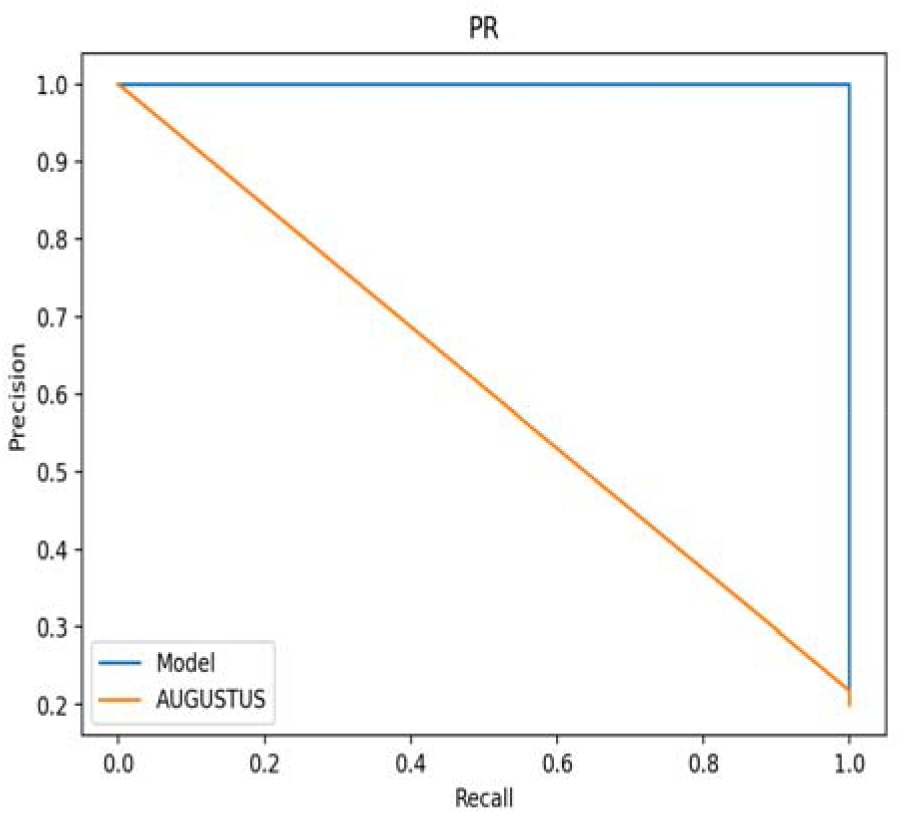
P-R curve for model and AUGUSTUS

**Figure 8.**
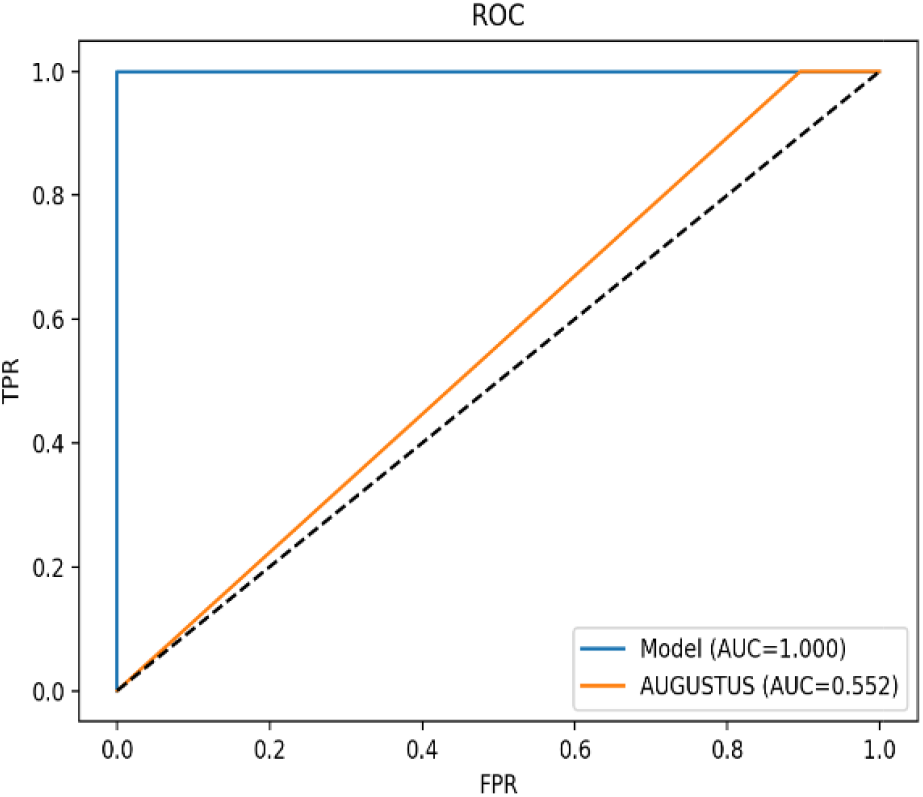
Comparison of model and AUGUSTUS

The ROC analysis reveals a substantial performance gap between the two approaches. The model achieves an AUC of 1.0, indicating perfect discrimination between coding and non-coding sequences across all thresholds. In contrast, AUGUSTUS attains an AUC of only 0.552, barely above random performance. Because AUGUSTUS produces binary predictions. These results underscore the advantages of probabilistic sequence models, which capture fine-grained coding signals that classical HMM-based gene finders fail to detect

## 6. Conclusion

In this work, we have presented an approach for building prediction functions of genes based on convolutional neural networks from real human genome sequences. The resulting model presents performances with average values of 98%, which compared to the works studied in the literature review on the same topic places it above them and positions it at the state of the art in developments for gene prediction based on machine learning, and in particular based on neural networks. Unlike our model, AUGUSTUS showed limited sensitivity to several perturbation types, particularly small indels and low-rate point mutations. This reflects the structure of its generalized HMM, which models coding potential using Markov chains and exon length distributions rather than frame consistency or fine-grained sequence features. As a result, perturbations that disrupt higher-order coding structure (e.g., codon-preserving shuffles) were more likely to flip AUGUSTUS predictions than biologically disruptive but statistically subtle mutations.

Taking into account the size of the training space (all coded sequences of the human genome), in an approach divide and conquer, we divided each chromosome into partitions containing sequences (examples) labeled with the name of the gene to which they belong. To construct the training set, we then implemented a novel method based on tf×idf matrices from the DNA sequences converted to amino acids sequences to identify and extract features for the creation of training examples. We evaluated the quality of the prediction of the induced functions using sequences of genes related to diseases caused by single-gene mutations. The excellent results confirm its potential use in medical applications and genomic research.

## 7. Future Research

In order to further improve the predictive capacity of our models, we have set a goal for the near future to develop algorithms for concurrent prediction (Ensemble Learning), combining deep learning CNN and Markovian methods as well as methods based on conditional probabilities.

## Code and Data Availability

Publicly available genomic datasets used in this study can be accessed from their respective repositories as cited in the manuscript. The preprocessing steps required to create the training and evaluation sets as well as the model creation are fully described in sections 3 (Design and Construction of the Model) and 4 (Training and Test Set selection). The core model architecture and training configuration are documented in sufficient detail to enable independent implementation.

The implementation of the proposed DNA sequences prediction method approach is part of an ongoing intellectual property evaluation. To maintain the integrity of this process, the full source code for the encoding module is not publicly released at this time. However, a high-level algorithmic description and all parameters necessaries for scientific assessment are provided in the manuscript and Appendix. Additional clarifications needed for reproducibility will be made available to qualified researchers upon reasonable request and under a non-commercial research agreement.

## Authors

Jesus Antonio Motta holds a Ph.D. in Computer Science with specialization in Artificial Intelligence. He has extensive experience as a systems engineer and university professor. His teaching and academic work span probability, linear and dynamic programming, stochastic processes, machine learning and bioinformatics. He has conducted research in natural language processing, image processing, and information extraction. His current research focuses on the development of bioinformatics tools and the analysis and design of intelligent systems

Pedro David Gomez is a physician specializing in Internal Medicine and a professor in this discipline. His academic and clinical work focuses on the diagnosis and management of complex medical conditions. His current professional activities center on advancing medical education and contributing to improved clinical outcomes through research and practice that include the study of autoimmune diseases and their relationship with genetic mutations.

