## Supplemental File 2 for "A Convolutional Deep Learning Approach to identify DNA Sequences for Gene Prediction"

Supplementary Material No. 2 for “**A Convolutional Deep Learning Approach to identify DNA Sequences for Gene Prediction”** article

This supplement contains the number of examples of each chromosome partition for the creation of the TS’s

TABLE S2_1

NUMBER OF EXAMPLES BY CHROMOSOME PARTITION

No. Chr. Part	Exs. No. Chr. Part. Exs. No. Chr. Part. Exs. No. Chr. Part. Exs.

| 1 | 4 | 4_0 | 5905 |
| --- | --- | --- | --- |
| 2 | 4 | 4_1 | 5551 |
| 3 | 4 | 4_3 | 4114 |
| 4 | 4 | 4_4 | 3193 |
| 5 | 4 | 4_5 | 4072 |
| 6 | 4 | 4_6 | 3942 |
| 7 | 6 | 6_0 | 4380 |
| 8 | 6 | 6_1 | 3982 |
| 9 | 4 | 4_2 | 5665 |
| 10 | 16 | 16_0 | 1595 |
| 11 | 16 | 16_1 | 2107 |
| 12 | 16 | 16_2 | 1656 |
| 13 | 16 | 16_3 | 1608 |
| 14 | 16 | 16_4 | 1638 |
| 15 | 16 | 16_5 | 1376 |
| 16 | 16 | 16_6 | 2323 |
| 17 | 16 | 16_7 | 1140 |
| 18 | 16 | 16_8 | 1514 |
| 19 | 16 | 16_9 | 2021 |
| 20 | 6 | 6_2 | 3165 |
| 21 | 6 | 6_3 | 3162 |
| 22 | 6 | 6_4 | 2923 |
| 23 | 6 | 6_5 | 4127 |
| 24 | 6 | 6_6 | 2464 |
| 25 | 6 | 6_7 | 1866 |
| 26 | 6 | 6_8 | 3365 |
| 27 | 6 | 6_9 | 2932 |
| 28 | X | X_0 | 3895 |
| 29 | X | X_1 | 4537 |
| 30 | X | X_2 | 3392 |
| 31 | X | X_3 | 3286 |
| 32 | X | X_4 | 2450 |
| 33 | X | X_5 | 2884 |
| 34 | X | X_6 | 2975 |
| 35 | 1 | 1_0 | 5189 |
| 36 | 1 | 1_1 | 4481 |
| 37 | 1 | 1_2 | 4782 |
| 38 | 1 | 1_3 | 3719 |
| 39 | 1 | 1_4 | 4611 |
| 40 | 1 | 1_5 | 4181 |
| 41 | 1 | 1_6 | 4232 |
| 42 | 1 | 1_7 | 4764 |
| 43 | 1 | 1_8 | 3527 |
| 44 | 1 | 1_9 | 2932 |
| 45 | 1 | 1_10 | 4106 |
| 46 | 1 | 1_11 | 4214 |
| 47 | 17 | 17_0 | 2471 |
| 48 | 17 | 17_1 | 1705 |
| 49 | 17 | 17_2 | 1786 |

| 50 | 17 | 17_3 | 1488 |
| --- | --- | --- | --- |
| 51 | 17 | 17_4 | 1637 |
| 52 | 17 | 17_5 | 1652 |
| 53 | 17 | 17_6 | 2019 |
| 54 | 13 | 13_0 | 2961 |
| 55 | 13 | 13_1 | 3469 |
| 56 | 13 | 13_2 | 2359 |
| 57 | 13 | 13_3 | 3865 |
| 58 | 13 | 13_4 | 1480 |
| 59 | 13 | 13_5 | 2482 |
| 60 | 19 | 19_0 | 1859 |
| 61 | 19 | 19_1 | 1755 |
| 62 | 19 | 19_2 | 1874 |
| 63 | 19 | 19_3 | 1955 |
| 64 | 19 | 19_4 | 1879 |
| 65 | 19 | 19_5 | 1839 |
| 66 | 19 | 19_6 | 1745 |
| 67 | 19 | 19_7 | 1913 |
| 68 | 19 | 19_8 | 1885 |
| 69 | 19 | 19_9 | 1830 |
| 70 | 12 | 12_0 | 4814 |
| 71 | 12 | 12_1 | 3625 |
| 72 | 12 | 12_2 | 3376 |
| 73 | 12 | 12_3 | 4528 |
| 74 | 12 | 12_4 | 4559 |
| 75 | 12 | 12_5 | 2716 |
| 76 | 12 | 12_6 | 4795 |
| 77 | 7 | 7_0 | 7137 |
| 78 | 7 | 7_1 | 5236 |
| 79 | 7 | 7_2 | 5248 |
| 80 | 7 | 7_3 | 3751 |
| 81 | 7 | 7_4 | 3387 |
| 82 | 7 | 7_5 | 4594 |
| 83 | 7 | 7_6 | 3155 |
| 84 | 11 | 11_0 | 3634 |
| 85 | 11 | 11_1 | 4760 |
| 86 | 11 | 11_2 | 3626 |
| 87 | 11 | 11_3 | 4189 |
| 88 | 11 | 11_4 | 4067 |
| 89 | 11 | 11_5 | 2652 |
| 90 | 11 | 11_6 | 3427 |
| 91 | 11 | 11_7 | 3185 |
| 92 | 18 | 18_0 | 2431 |
| 93 | 18 | 18_1 | 1679 |
| 94 | 18 | 18_2 | 1577 |
| 95 | 18 | 18_3 | 1736 |
| 96 | 18 | 18_4 | 426 |
| 97 | 18 | 18_5 | 1525 |
| 98 | 21 | 21_0 | 1186 |

| 99 | 21 | 21_1 | 969 |
| --- | --- | --- | --- |
| 100 | 21 | 21_2 | 276 |
| 101 | 21 | 21_3 | 1117 |
| 102 | 21 | 21_4 | 193 |
| 103 | 21 | 21_5 | 1002 |
| 104 | 5 | 5_0 | 4874 |
| 105 | 5 | 5_1 | 3664 |
| 106 | 5 | 5_2 | 4614 |
| 107 | 5 | 5_3 | 3994 |
| 108 | 5 | 5_4 | 3613 |
| 109 | 5 | 5_5 | 1530 |
| 110 | 5 | 5_6 | 3251 |
| 111 | 8 | 8_0 | 4292 |
| 112 | 8 | 8_1 | 2188 |
| 113 | 8 | 8_2 | 3166 |
| 114 | 8 | 8_3 | 3190 |
| 115 | 8 | 8_4 | 2033 |
| 116 | 8 | 8_5 | 1872 |
| 117 | 8 | 8_6 | 3493 |
| 118 | Y | Y_0 | 555 |
| 119 | Y | Y_1 | 632 |
| 120 | Y | Y_2 | 914 |
| 121 | Y | Y_3 | 555 |
| 122 | Y | Y_4 | 541 |
| 123 | Y | Y_5 | 742 |
| 124 | 2 | 2_0 | 3786 |
| 125 | 2 | 2_1 | 3016 |
| 126 | 2 | 2_2 | 3755 |
| 127 | 2 | 2_3 | 1535 |
| 128 | 2 | 2_4 | 3195 |
| 129 | 2 | 2_5 | 4003 |
| 130 | 2 | 2_6 | 2613 |
| 131 | 2 | 2_7 | 3315 |
| 132 | 2 | 2_8 | 1480 |
| 133 | 2 | 2_9 | 772 |
| 134 | 2 | 2_10 | 3322 |
| 135 | 2 | 2_11 | 3680 |
| 136 | 10 | 10_0 | 2109 |
| 137 | 10 | 10_1 | 2416 |
| 138 | 10 | 10_2 | 1248 |
| 139 | 10 | 10_3 | 1499 |
| 140 | 10 | 10_4 | 1962 |
| 141 | 10 | 10_5 | 1894 |
| 142 | 10 | 10_6 | 1304 |
| 143 | 14 | 14_0 | 966 |
| 144 | 14 | 14_1 | 1041 |
| 145 | 14 | 14_2 | 1462 |
| 146 | 14 | 14_3 | 209 |
| 147 | 14 | 14_4 | 657 |

| 148 | 14 | 14_5 | 1960 |
| --- | --- | --- | --- |
| 149 | 14 | 14_6 | 1827 |
| 150 | 9 | 9_0 | 1574 |
| 151 | 9 | 9_1 | 1449 |
| 152 | 9 | 9_2 | 1233 |
| 153 | 9 | 9_3 | 1515 |
| 154 | 9 | 9_4 | 790 |
| 155 | 9 | 9_5 | 1698 |
| 156 | 9 | 9_6 | 1015 |
| 157 | 9 | 9_7 | 1806 |
| 158 | 15 | 15_0 | 3384 |
| 159 | 15 | 15_1 | 3085 |
| 160 | 15 | 15_2 | 3063 |
| 161 | 15 | 15_3 | 3277 |
| 162 | 15 | 15_4 | 2552 |
| 163 | 15 | 15_5 | 3391 |
| 164 | 20 | 20_0 | 569 |
| 165 | 20 | 20_1 | 644 |
| 166 | 20 | 20_2 | 600 |
| 167 | 20 | 20_3 | 403 |
| 168 | 20 | 20_4 | 709 |
| 169 | 20 | 20_5 | 1195 |
| 170 | 20 | 20_6 | 999 |
| 171 | 22 | 22_0 | 310 |
| 172 | 22 | 22_1 | 356 |
| 173 | 22 | 22_2 | 521 |
| 174 | 22 | 22_3 | 355 |
| 175 | 22 | 22_4 | 104 |
| 176 | 22 | 22_5 | 529 |
| 177 | 3 | 3_0 | 2836 |
| 178 | 3 | 3_1 | 3068 |
| 179 | 3 | 3_2 | 2727 |
| 180 | 3 | 3_3 | 2100 |
| 181 | 3 | 3_4 | 4087 |
| 182 | 3 | 3_5 | 2891 |
| 183 | 3 | 3_6 | 2728 |
| 184 | 3 | 3_7 | 2363 |
| 185 | 3 | 3_8 | 1161 |
| 186 | 3 | 3_9 | 1809 |
| 187 | 3 | 3_10 | 3625 |
| 188 | 3 | 3_11 | 2624 |
| 189 | Y | Y_6 | 573 |
| 190 | 17 | 17_7 | 1737 |
| 191 | 17 | 17_8 | 1302 |
| 192 | 17 | 17_9 | 1787 |
| 193 | 17 | 17_10 | 2084 |
| 194 | 17 | 17_11 | 1667 |

**Total number of examples: 491463**
