## Supplemental File 1 for "A Convolutional Deep Learning Approach to identify DNA Sequences for Gene Prediction"

Supplementary Material No. 1 for “**A Convolutional Deep Learning Approach to identify DNA Sequences for Gene Prediction”** article

This supplement contains the ROC curves of 16 genes of interest compared against 6 random gene ROC curves

| 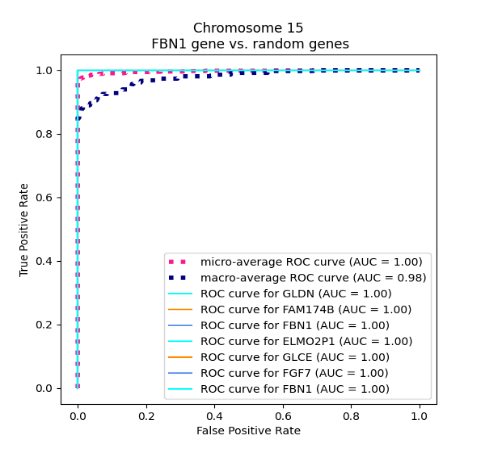 | 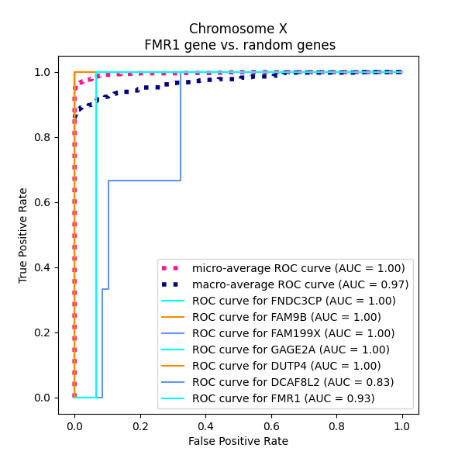 |
| --- | --- |
| 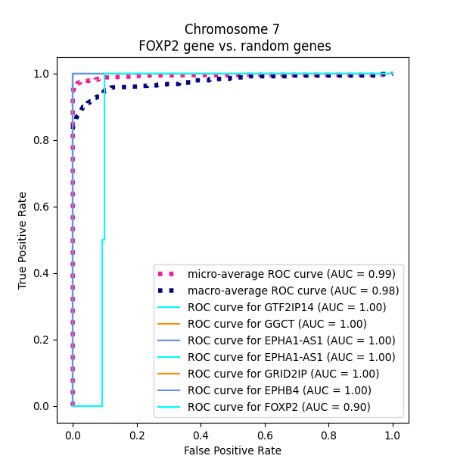 | 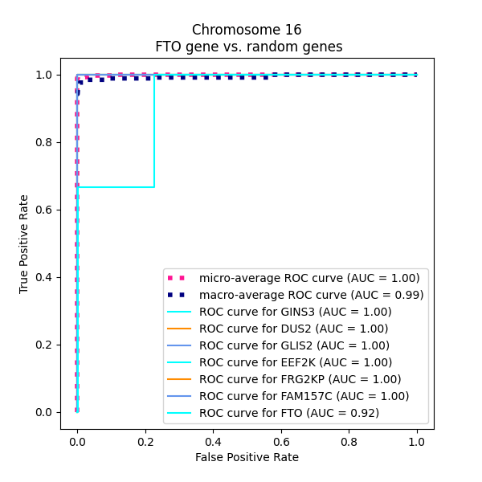 |

Fig. S1_1. Genes of Interest vs. random genes

| 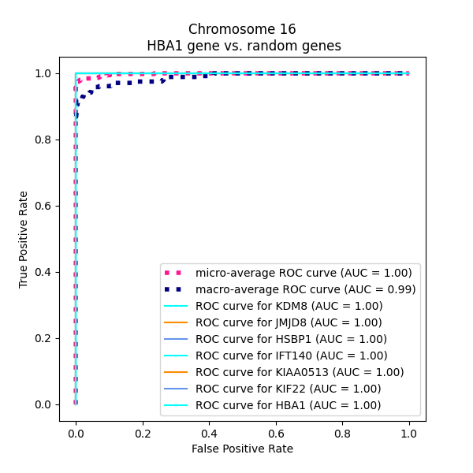 | 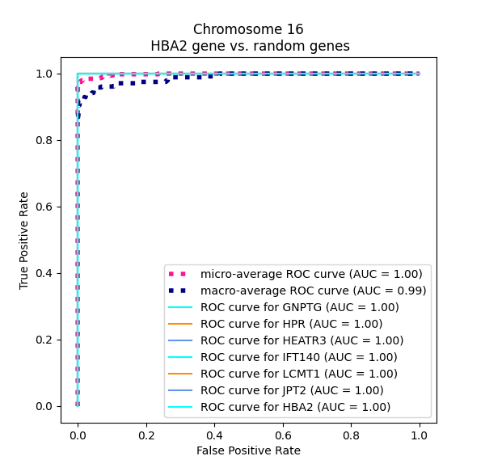 |
| --- | --- |
| 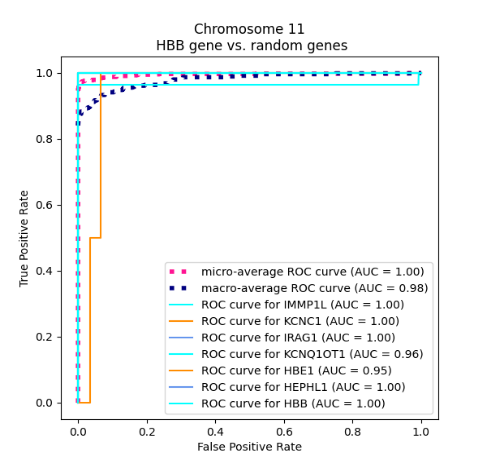 | 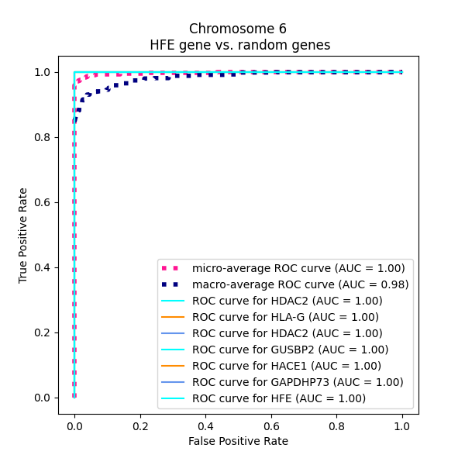 |

Fig. S1_2. Genes of Interest vs. random genes

| 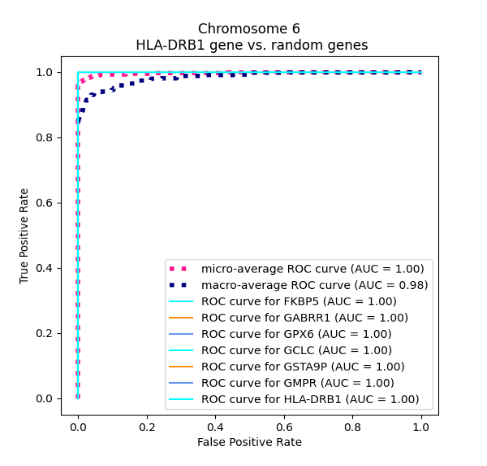 | 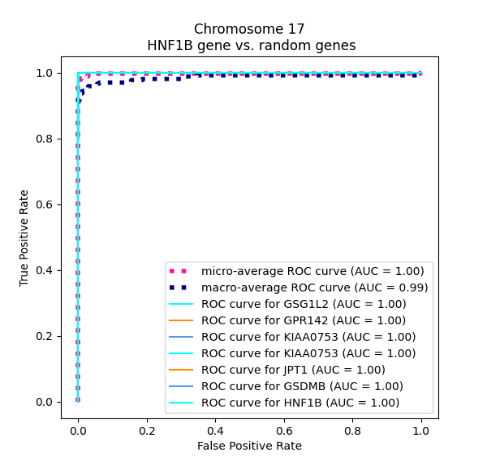 |
| --- | --- |
| 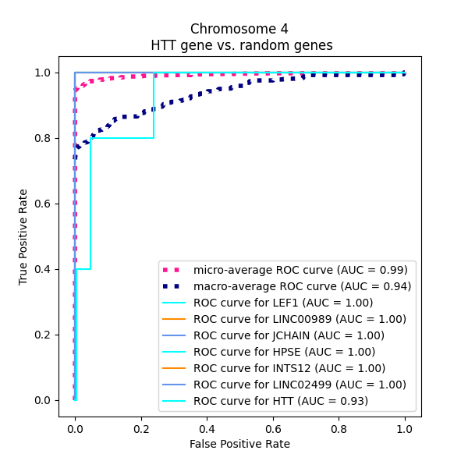 | 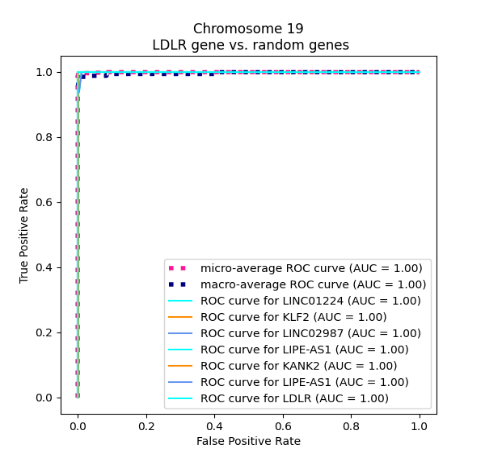 |

Fig. S1_3. Genes of Interest vs. random genes

| 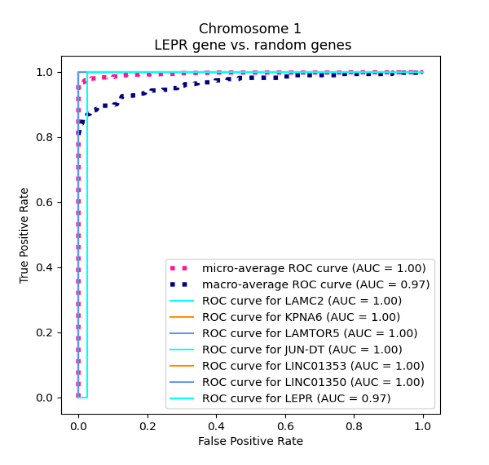 | 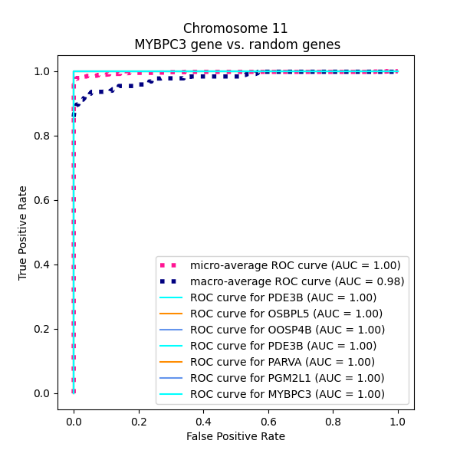 |
| --- | --- |
| 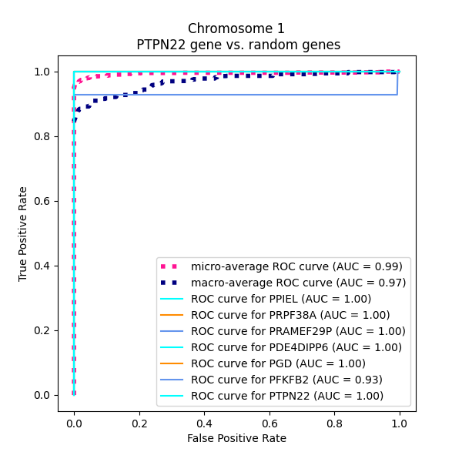 | 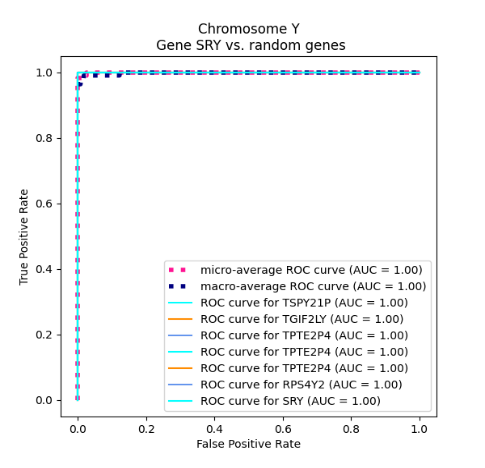 |

Fig. S1_4. Genes of Interest vs. random genes
